# Modulation of the microhomology-mediated end joining pathway suppresses large deletions and enhances homology-directed repair following CRISPR-Cas9-induced DNA breaks

**DOI:** 10.1101/2022.11.16.516713

**Authors:** Baolei Yuan, Chongwei Bi, Jinchen Wang, Yiqing Jin, Khaled Alsayegh, Muhammad Tehseen, Gang Yi, Samir Hamdan, Yanyi Huang, Mo Li

**Affiliations:** Bioscience Program, Biological and Environmental Sciences and Engineering Division, King Abdullah University of Science and Technology (KAUST), Thuwal 23955-6900, Kingdom of Saudi Arabia; Beijing Advanced Innovation Center for Genomics (ICG), Biomedical Pioneering Innovation Center (BIOPIC), School of Life Sciences, College of Chemistry, College of Engineering, Peking-Tsinghua Center for Life sciences, Peking University, Beijing, China; Institute for Cell Analysis, Shenzhen Bay laboratory, Shenzhen, China; Bioengineering Program, Biological and Environmental Sciences and Engineering Division, King Abdullah University of Science and Technology (KAUST), Thuwal 23955-6900, Kingdom of Saudi Arabia

## Abstract

CRISPR-Cas9, an efficient genome editing tool, has been widely used in research and holds great promise in the clinic. However, large unintended rearrangements of the genome occur frequently after CRISPR-Cas9 editing and their potential risk cannot be ignored. In this study, we detected large deletions (LDs) induced by CRISPR-Cas9 in human embryonic stem cells (hESCs) and found the microhomology end joining (MMEJ) DNA repair pathway plays a predominant role in LD. We genetically targeted PARP1, RPA, POLQ and LIG3, which play critical roles in MMEJ, during CRISPR-Cas9 editing. By analyzing LD events in two independent gene loci, CD9 and PIGA, using flow cytometry and long-read individual molecule sequencing (IDMseq), we showed that knocking down PARP1 and LIG3 does not alter the frequency of Cas9-induced LD, while knocking down or inhibiting POLQ dramatically reduces LD. Knocking down RPA increases LD frequency, and overexpression of RPAs consistently reduces LD frequency. Interestingly, small-molecule inhibition of POLQ and delivery of recombinant RPA proteins also dramatically increase the efficiency of homology-directed repair (HDR). In conclusion, RPA and POLQ play opposite roles in Cas9-induced LD, modulation of POLQ and RPA can reduce LD and improve HDR, thus holding promise for safe and precise genome editing.

## INTRODUCTION

CRISPR-Cas9 can introduce double strand breaks (DSBs) to a specific genomic locus that shares sequence complementarity with the CRISPR guide RNA (gRNA). The DSBs can be repaired through different cellular pathways, including classical non-homology end joining (C-NHEJ, hereafter referred to as NHEJ), MMEJ (also called alternative NHEJ), homologous recombination (HR) and single-stranded annealing (SSA). NHEJ often generates small insertions and deletions (indels) ^1^ and is believed to be the dominant repair pathway for DSB induced by CRIPSR-Cas9 ^2^. MMEJ relies on small homologies for DNA repair, while SSA requires longer ones. HR is an error-free DNA repair mechanism that needs a homologous template.

The majority of on-target indels induced by CRISPR-Cas9 are estimated to be less than 20 bp in numerous large-scale studies on Cas9 cleavage outcomes ^1,3-5^. However, on-target large deletions (LDs) and large complex rearrangements of the genome caused by CRISPR-Cas9 were not appreciated until recently ^6^. The reason is that short PCR amplicons (usually less than 300 bp) were used in previous studies to analyze the genome editing outcomes by Illumina or Sanger sequencing, which are not able to detect LDs or more complex genomic rearrangements. Long-read sequencing platforms, such as PacBio and Nanopore that are much better at resolving large rearrangements, have been used for the analysis of genome editing outcomes ^6-10^.

The LD issue can have significant implications for the application of the versatile genome editing tool CRISPR-Cas9. We found that the breakpoint junctions of LDs induced by CRISPR-Cas9 are highly enriched in microhomologies (MHs). We hypothesized that MMEJ play roles in generating LDs following CRISPR-Cas9 cleavage, and investigated the roles of four key MMEJ genes–Poly [ADP-ribose] polymerase 1 (PARP1), replication protein A (RPA), DNA polymerase theta (POLQ), and DNA ligase 3 (LIG3)–in LD formation. In brief, PARP1 binds the ends of the DSB as a competitor of Ku that is well known to play a similar role in the NHEJ pathway ^11^, thus channeling DNA repair to the MMEJ pathway. RPA that contains three subunits–RPA1, RPA2, and RPA3– binds resected single-stranded DNA (ssDNA) and triggers homologous recombination (HR) ^12,13^. RPA also prevents ssDNA annealing thus further blocking MMEJ repair ^14^. POLQ plays multiple roles in MMEJ that include promoting DNA synapse formation and ssDNA annealing, extending overhangs ^15^, and inhibiting HR ^16^, and therefore is considered to play a central role in MMEJ in higher organisms ^17^. LIG3 is a major ligase for sealing the gaps in the last step of MMEJ ^17,18^.

We found that depletion or inhibition of POLQ or overexpression of RPA significantly reduced LD frequency. By contrast, knocking down PARP1 or LIG3 had no effect on LD frequency. We further provided base-resolution validation of the observations by using a long-read individual molecular sequencing method, IDMseq ^7^. We also found small-molecule inhibition of POLQ or delivery of recombinant RPA proteins significantly increased HDR efficiency. These results highlight the role of MMEJ in CRISPR-Cas9 mediated genome editing and provide potential targets and strategies for safe and precise genome editing.

## RESULTS

### Most CRISPR-Cas9 induced LDs in human pluripotent stem cells contain microhomology at breakpoint junctions

CRISPR-Cas9 can efficiently cut target DNA to promote gene knockout through the formation of small indels or precise installation of DNA sequence changes through homology directed repair (HDR). However, it also causes unintended LDs and structural variations (SVs) up to several megabases or even whole chromosome loss ^6,7,19-21^. The underlying mechanism of CRISPR-Cas9 induced LD remains unclear. We collated sequencing data of 329 CRISPR-Cas9 edited alleles from two published studies ^6,19^, and found an unusually high frequency of MHs at LD breakpoint junctions. For example, MHs ≥ 2 bp were present in more than 70% of LD alleles (Fig. 1a). We also randomly examined four clones of human pluripotent stem cell (hPSC) lines edited by CRISPR-Cas9 in-house and found one harbored a LD that contained MH (Fig. 1b, Fig. S1a).

**Fig. 1.**
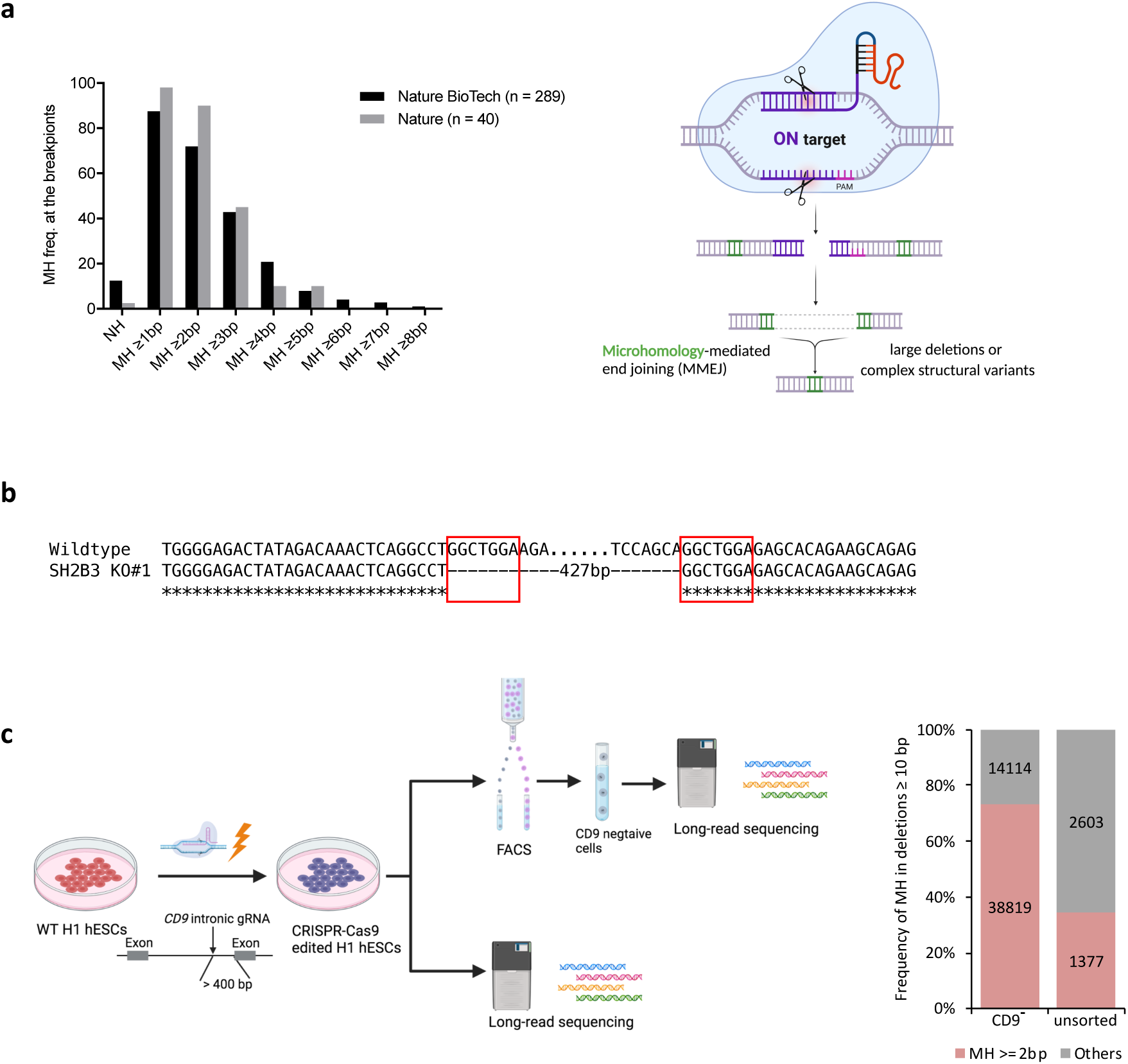
MH is enriched at the breakpoint junctions of CRISPR-Cas9 induced LDs. **a** Left: analysis of microhomology (MH) frequency at Cas9 induced breakpoint junctions in two published data ^6,19^; right: schematic of how MMEJ could lead to CRISPR-induced LDs (created with BioRender.com). NH: no homology **b** LD event detected in a Cas9-edited human pluripotent cell line. Red color boxes indicate MH sequences. **c** Left: schematic of the strategy to analyze CRISPR-induced LDs in the *CD9* locus; right: MH frequency in deletions ≥ 10 bp quantified from long-read sequencing data.

To provide quantitative evidence for prevalent MHs in Cas9-edited loci, we edited the *CD9* and *PIGA* intronic regions, respectively, in H1 hESCs using Cas9/gRNA ribonucleoprotein (RNP) complex (Fig. 1c, Fig. s1b). In both loci, the distance between the intronic gRNA and the nearest exons is more than 200 bp. Therefore, the edited cells that lose cell surface expression of *CD9* or *PIGA*, as monitored by fluorescence-activated cell sorting (FACS), are considered to contain LDs that extend at least from the CRISPR-Cas9 cleavage site to the nearest exon. We performed PacBio circular consensus sequencing of a 7-kb region flanking the intronic gRNA target amplified from the CD9 or PIGA negative cells and respective unsorted cells (Fig. 1c, Fig. s1b). The analysis of reads with deletions ≥ 10-bp showed that 80.2% and 73.3% of breakpoint junctions in the *CD9* and *PIGA* target sites, respectively, contained MHs ≥ 2 bp in negative populations, and the MH frequency dropped in both unsorted populations (Fig. 1c, Fig. s1b). Since MMEJ mediated DNA repair results in MH at the breakpoint junctions (Fig. 1a), the high occurrence of MHs suggest that the MMEJ pathway plays roles in meditating CRISPR-induced LDs.

### RPA and POLQ but not PARP1 and LIG3 play roles in LD formation

To better understand the role of the MMEJ pathway in the formation of LD, we modulated the function of four genes (PARP1, RPA, POLQ, and LIG3) in hPSCs undergoing CRISPR-Cas9 editing (Fig. 2a, b). To achieve consistent and uniform induction of CRISPR-Cas9 editing, we used an H1 ESC line with a doxycycline-inducible Cas9 expression system knocked-in to the AAVS1 safe harbor locus (H1-iCas9) ^22^. We firstly investigated the range of LD frequency by using sgRNAs targeting different intronic regions of the X-linked PIGA gene, which is an established model ^3,6,23^ for the study of CRISPR-Cas9 editing outcomes (sgRNA positions are shown in Fig. s1c). Thirteen intronic sgRNAs were individually expressed in H1-iCas9 ESCs using a constitutive lentiviral vector. Upon doxycycline induction, these sgRNAs guided Cas9 to generate DSBs located 126-489 bp from the nearest exon (Fig. s1c). Subsequent DNA repair could lead to small indels that did not reach coding sequences and preserved PIGA expression, or, to LDs that extended into nearby exons and disrupted PIGA expression, which resulted in cells stained positively and negatively with FLAER reagent, respectively (Fig. s1c). Control sgRNAs that target exons led to a nearly 100% PIGA knockout based on the FLAER FACS assay, suggesting that our system reached a saturating level of editing (Fig. s1c).

**Fig. 2.**
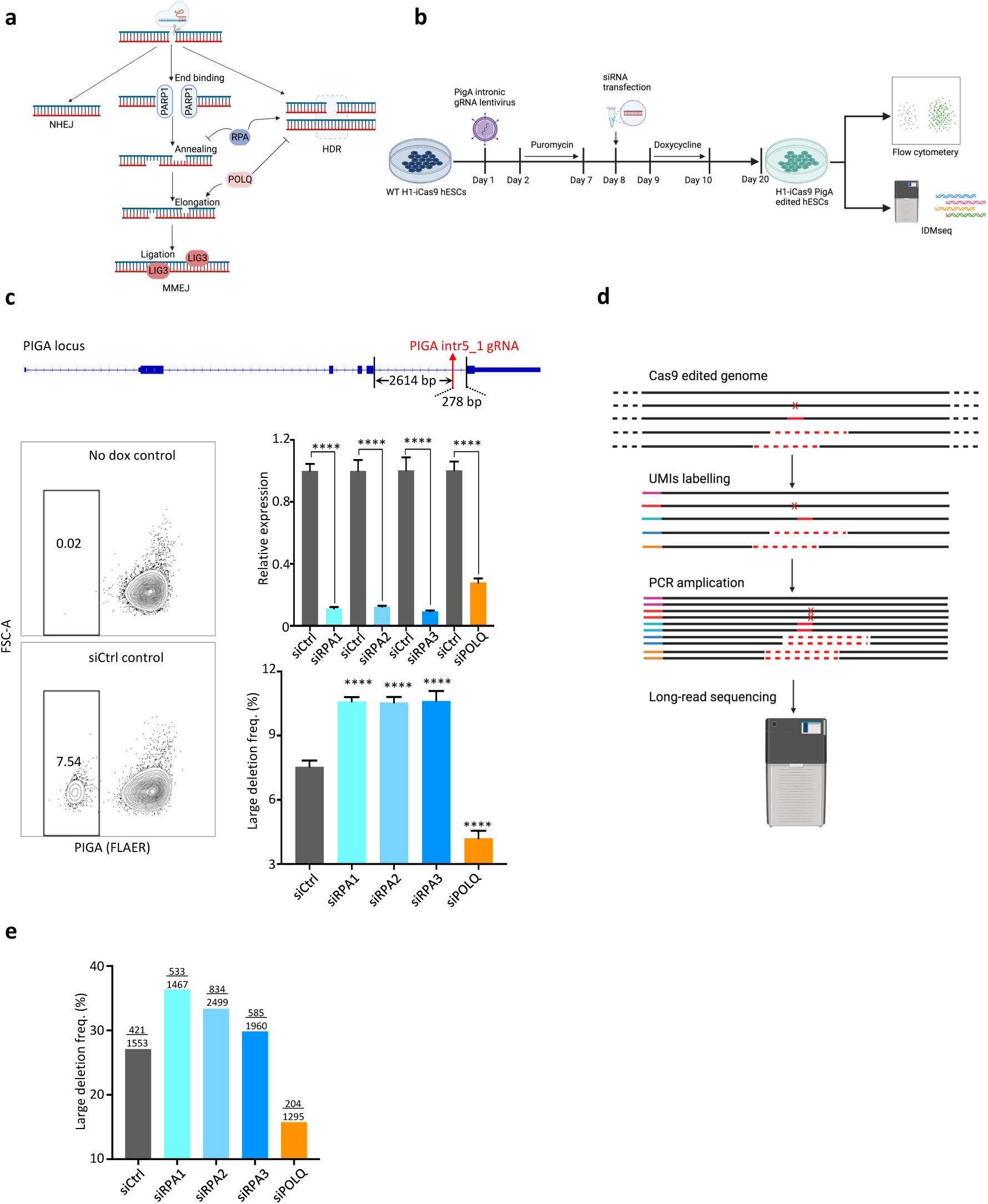
RPA and POlQ play opposite roles in CRISPR-Cas9 induced LDs. **a** Schematic of the roles of four key genes in the MMEJ pathway (created with BioRender.com). **b** Diagram of the workflow for the knockdown experiments (created with BioRender.com). **c** Top: the location of the PIGA intronic gRNA; bottom left: example flow cytometry analysis of PIGA expression using the FLAER assay; bottom right: normalized mRNA level of siRNA target genes and LD frequency quantified by FACS, biological replicates *n* =3, **** *p* < 0.0001. **d** Schematic of IDMseq analysis of LDs. **e** Frequency of LD (≥ 30 bp) quantified by IDMseq. The nominator indicates the LD event number, and the denominator indicates the total event number detected by IDMseq.

The intronic sgRNA data showed that the frequency of PIGA deficient cells (FLAER^neg^) ranged from 0.23% to 9.05% (average 3.52±0.83%) in Cas9-edited cells (Fig. s1c). Note that FLAER^neg^% is likely a conservative estimate of LD%, because LDs extending to the opposite side of the nearest exon may not result in PIGA deficiency (i.e., FLAER^neg^) and in-frame LDs may lead to hypomorpic PIGA levels (cells with intermediate FLAER staining in Fig. s1c). Background LD events in cells without Cas9 expression (no dox) were nearly undetectable (Fig. s2a), thus proving that the LDs were due specifically to Cas9 induced DSBs. LD frequency could not be predicted solely by the orientation of the sgRNA (targeting the + or – strand) or the distance between the sgRNA and the nearest exon (Fig. s1c, d), and could be sequence context dependent. We selected intr5_1 sgRNA, located 278 bp away from the nearest exon, that induced the highest LD% (Fig. s1c) as a sensitive setup to evaluate the effects of manipulating the MMEJ pathway on LD frequency in the rest of our study.

Considering the prevalence of MHs in LDs, we hypothesized that LD frequency could be controlled by modulating the activity of the MMEJ pathway. We knocked down four key players of the MMEJ pathway, PARP1, LIG3, RPA (including RPA1, RPA2, and RPA3), and POLQ in H1-iCas9 cells expressing the PIGA intr5_1 sgRNA (Fig. 2a-c, S2b) and induced Cas9 expression 24 hours later. LD frequency was monitored by FACS analysis of FLAER staining as described in the preceding paragraph (Fig. 2b, c; Fig. s2a). The results showed that knocking down POLQ caused a 40% reduction in LD frequency, while knocking down RPA proteins lead to a 40% increase in LD frequency (Fig. 2c). Knocking down PARP1 or LIG3 consistently showed no effect on LD frequency (Fig. s2b), so we excluded them from further study.

To better quantify the Cas9 editing outcomes in MMEJ-knockdown H1-iCas9 cells at base resolution, we next performed IDMseq ^7^ of the *PIGA* locus. Briefly, individual genomic regions flanking the Cas9 cut site were labelled with a unique molecular identifier (UMI) and amplified for long-read PacBio sequencing (Fig. 2d). In the subsequence sequencing data analysis, we referred to deletions ≥ 30 bp as LDs. The baseline LD frequency of the control siRNA as per IDMseq was higher than that estimated by FACS, which is expected because FLAER^neg^% underestimates LD frequency as discussed above and because deletions 30-278 bp in size are only detectable by IDMseq. Consistent with the FACS analysis the IDMseq results showed that knocking down POLQ and RPA decreased and increased LD frequency, respectively (Fig. 2e).

### LD induced by CRIPSR-Cas9 can be controlled by modulating POLQ and RPA

POLQ is an error-prone polymerase and is upregulated in numerous cancers ^24-28^. The antibiotic novobiocin (NVB) has recently been identified as a specific inhibitor of POLQ. NVB inhibits the ATPase activity of POLQ through direct binding to its ATPase domain and thus phenocopies POLQ depletion and impairs MMEJ DNA repair in human cells ^26^. We therefore used NVB to test if targeting a specific MMEJ-related activity of POLQ could recapitulate the reduction of LD frequency through knocking down POLQ. NVB was introduced to the cells during the doxycycline induction of Cas9 expression in the intr5_1 sgRNA-positive H1-iCas9 ESCs. NVB reduced LD frequency up to 50% in a dose dependent manner (Fig. 3a). No effect of NVB on pluripotency was detected in treated hESCs (Fig. s2c). High concentrations of NVB (50 µM) showed signs of cytotoxicity, while lower concentrations were well tolerated (Fig. s2d). These results showed that transient inhibition of POLQ activity is sufficient to reduce the formation of LDs following repair of DSBs induced by Cas9.

**Fig. 3.**
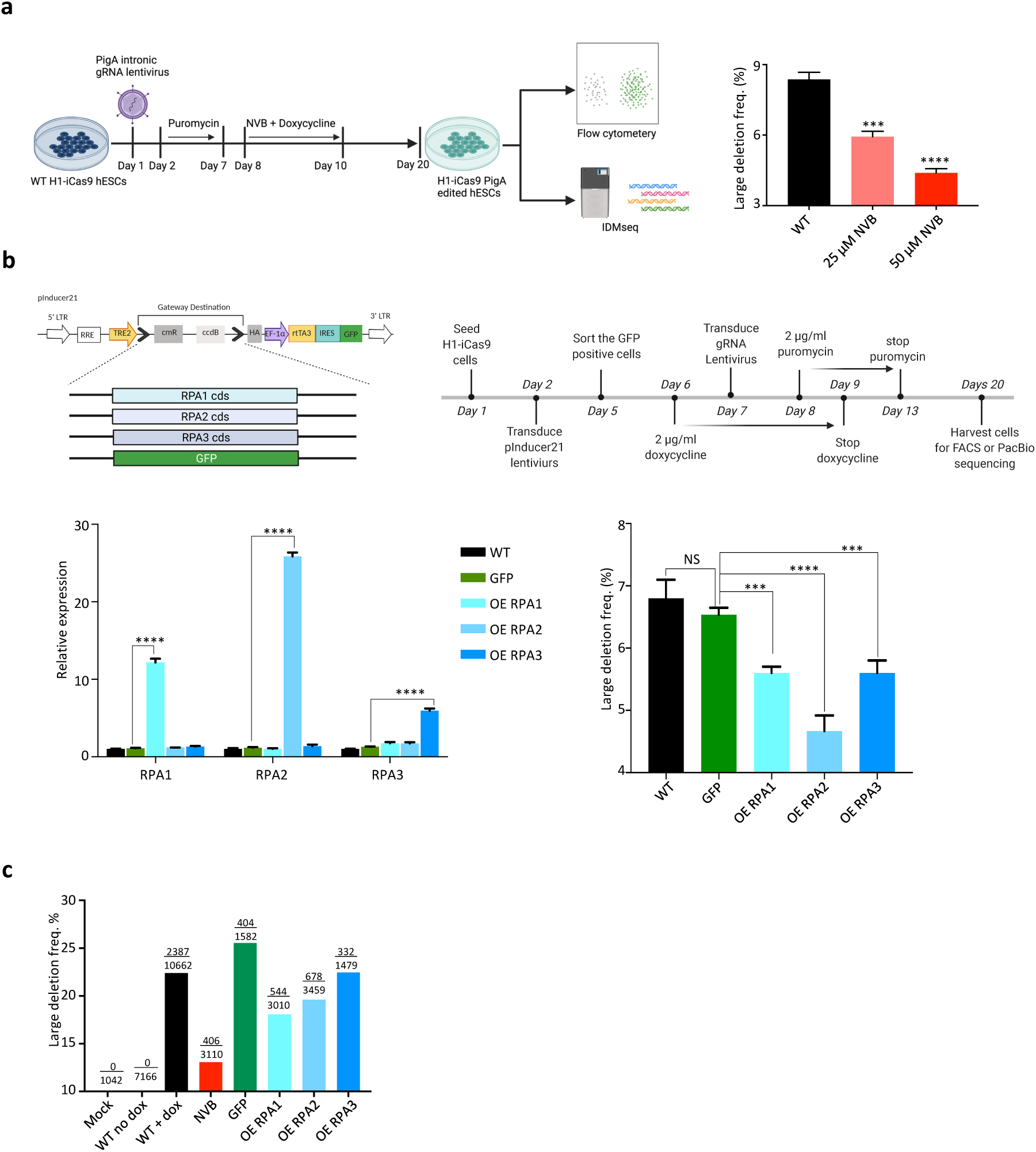
CRISPR-Cas9-induced LDs can be suppressed by inhibiting POLQ or overexpressing RPA. **a** Left: schematic of the workflow for the POLQ inhibition experiment (created with BioRender.com); right: LD frequency quantified by FACS, *** *p* < 0.001, **** *p* < 0.0001. **b** Top left: Schematics of the inducible RPA constructs; top right: the workflow for RPA overexpression experiments; bottom left: the mRNA level of RPA genes after doxycycline treatment for two days; bottom right: LD frequency quantified by FACS, biological replicates *n* =3, *** *p* < 0.001, **** *p* < 0.0001. OE: overexpression. **c** Frequency of LD (≥ 30 bp) quantified by IDMseq. The nominator indicates the LD event number, and the denominator indicates the total event number detected by IDMseq.

The RPA proteins prevent ssDNA annealing, thus further blocking MMEJ repair ^14,29^. Consistently, we showed that knocking down RPA increases LD. We hypothesized that increasing RPA availability could divert DNA repair away from the MMEJ pathway during Cas9 editing and lead to a reduction of LD. Therefore, we cloned three RPA subunits (RPA1, RPA2, RPA3) and GFP (as a control) individually into an inducible lentiviral expression vector, pInducer21, that expresses GFP constitutively (Fig. 3b). Successfully transduced cells were sorted based on GFP positivity and transgene expression was induced by doxycycline. The expression level of the transgenic RPA proteins increased between 6-fold to 26-fold after doxycycline induction without affecting the expression of other RPA subunits (Fig. 3b). Such levels of overexpression of all three RPA proteins resulted in significant reductions in LD frequency as detected by FACS (Fig. 3b).

To gain a sequence-level understanding of the effect of POLQ inhibition and RPA overexpression on DNA repair outcome of CRISPR-Cas9 editing, we performed IDMseq ^7^ of the PIGA locus as in the knockdown experiments. The results are consistent with the FACS analysis. Both POLQ inhibition and RPA overexpression can reduce CRISPR-Cas9-induced LD (Fig. 3c).

To examine whether modulating POLQ activity or RPA overexpression affected the desirable small indel formation, we analyzed the editing efficiency of an sgRNA targeting PIGA exon 2 by FACS (Fig. s2e). The data showed that treatment with NVB or overexpression of RPA1, RPA2, and RPA3 did not change the frequency of PIGA knockout cells (majority of which contain small indels). These results showed that LD can be controlled by modulating POLQ activity and overexpression of RPA without changing the overall editing efficiency. Together, there data suggested that small-molecule inhibition of POLQ and RPA overexpression could offer convenient and safe ways to reduce unwanted LDs following Cas9 editing without compromising editing efficiency.

### Modulation of POLQ and RPA can enhance HDR efficiency

The inhibition of the MMEJ pathway could potentially increase HDR efficiency. To test this hypothesis, we established a GFP mutant hPSC line that can be repaired via HDR by delivering Cas9/sgRNA RNP and an ssODN donor (Fig. 4a). Compared to the control, 25 µM NVB treatment for 24 hours before gene editing significantly improved the HDR efficiency. However, only low-dose (less than 10 pmol) recombinant RPA, premixed with Cas9/sgRNA RNP and ssODN donor before electroporation, resulted in increased HDR efficiency (Fig. 4b; Fig. s3a-c). The combination of NVB treatment and low-dose RPA further improved HDR efficiency (Fig. 4b; Fig. s3b, c). To understand why high-dose recombinant RPA reduced HDR efficiency, we monitored the delivery of 5’ Cy3-labelled ssODN in the cells using FACS after co-electroporation with Cas9/sgRNA RNP and different doses of RPA. The results showed that recombinant RPA negatively affected ssODN delivery in a dose-dependent manner, and low-dose RPA (2.5 pmol or less) did not result in any obvious reduction of ssODN (Fig. s3d). Hence, modulation of POLQ activity and RPA level can enhance HDR efficiency.

**Fig. 4.**
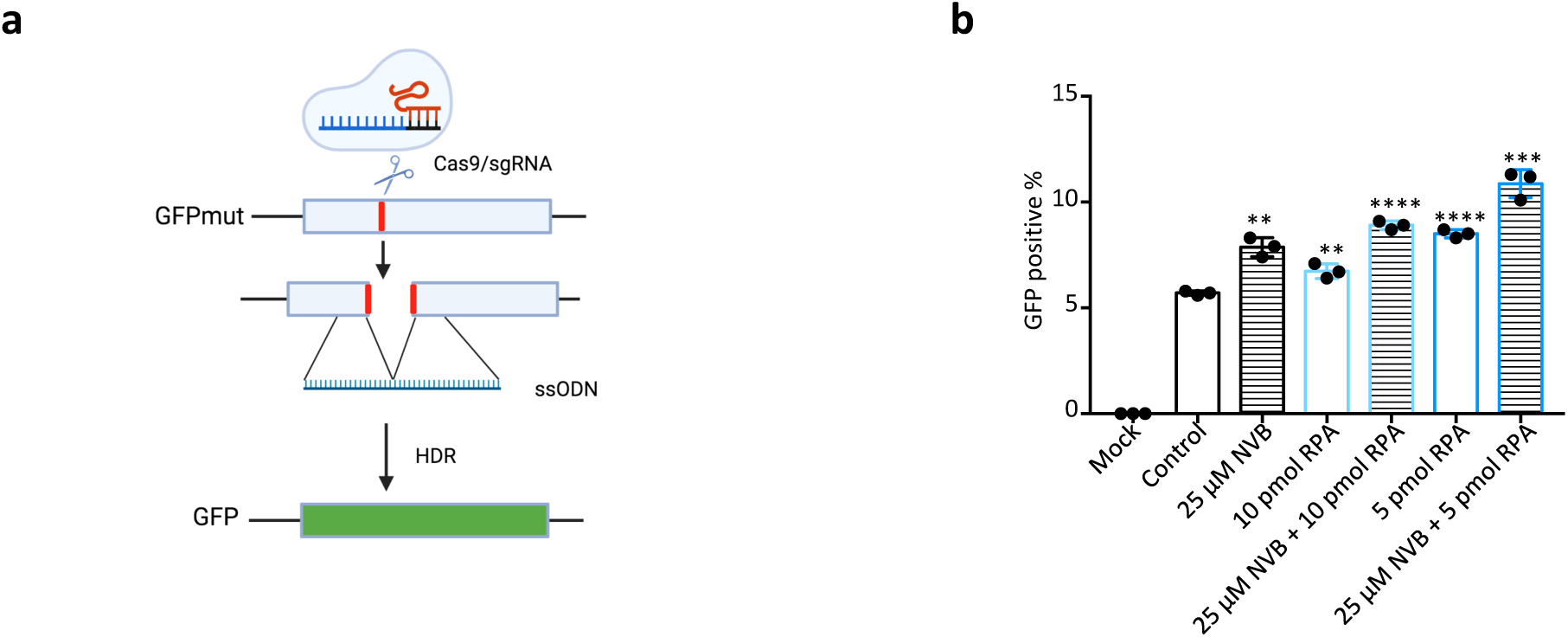
Modulation of POLQ and RPA increases HDR efficiency. **a** Schematic of mutant GFP correction by Cas9/sgRNA and ssODN via HDR. The green color indicates the restoration of green fluorescence. **b** The frequency of GFP positive cells quantified by FACS, biological replicates *n* =3, ** p < 0.01, *** *p* < 0.001, **** *p* < 0.0001.

## DISCUSSION

Our study showed a high frequency of MHs at Cas9-induced LD breakpoint junctions. This phenomenon was also reported when a dual paired Cas9^D10A^ nickases or a paired Cas9 nuclease system was used in genome editing ^30^ and in a study on Cas9 editing in mouse embryonic stem cells ^23^. Increasing evidence proved that MMEJ is not only a backup repair pathway but also an active one when HR and NHEJ are intact ^17,31^. That CRISPR-Cas9 continuously recuts the target after error-free DNA repair (which regenerates the target) could increase the chance for LD-prone repair pathway MMEJ. Although how asymmetrical release of the 3’ end of non-target DNA strand after Cas9 cleavage and long-term residence of Cas9 on the broken ends of DNA ^32^ affect DNA repair pathway choice is unclear, that knocking down PARP1 does not affect LD frequency suggests CRISPR-Cas9 induced DSBs initiate MMEJ repair pathway via a PARP1-independent manner (Fig. s2b). Although LIG3 is a predominant ligase of the MMEJ pathway that seals the nicks in DNA, its function could be replaced by other ligases, for example, LIG1 ^16^. The deficiency of PARP1 and LIG3 also did not affect LD in Cas9-edited mouse ESCs ^23^. POLQ plays a central role in MMEJ in higher organisms ^17^. Knocking down or inhibiting POLQ caused significant reduction of LD, which suggested limited functional redundancy between POLQ and other DNA polymerases and reaffirmed the central role of MMEJ in Cas9-induced LD. RPA plays important roles in DNA replication and DNA repair. Lack of RPA leads to more frequent LDs induced by CRISPR-Cas9. The reason for this may that without RPA binding the naked ssDNA resected from DSBs could have a higher chance of annealing at MHs.

An increasing number of studies have been conducted on CRISPR-Cas9 induced LDs. One study showed that LD can be prevented by enhanced homology-directed repair (HDR) via delivering ssODNs or adeno-associated virus (AAV) donors and NHEJ mediated dsODN insertion in primary T cells and hematopoietic stem cells but not in iPSCs ^8^. It can be construed as evidence that suppression of MMEJ via promoting the HDR and NHEJ pathways could reduce LD, which complement our direct mechanistic insights into MMEJ. Another study targeted 32 DNA repair genes associated with the NHEJ, MMEJ and HR repair pathways in mouse ESCs, and found that NHEJ pathway hindered LD while MMEJ pathway promoted LD ^33^, which is consistent with our results. RPA, a key player in LD discovered in our study, was not included in the 32 genes. A recent study showed that LDs and translocations can be reduced in T cells by using a Cas9 fused with an optimized exonuclease TREX2, which prevents perfect DNA repair ^34^. Although these tools are promising, they did not provide new insights to understand LD. CRISPR-Cas engineering strategies such as base editor and primer editor do not induce DSBs and can potentially perform precise editing without unwanted LD ^10^.

In this study, we first discovered that two key MMEJ genes (POLQ and RPA) regulate CRISPR-Cas9 induced LD formation and provided a mechanistic understanding of Cas9-induced LD. We then demonstrated that small-molecule inhibition of POLQ or supplying recombinant RPA together with Cas9/sgRNA RNP and ssODN can significantly reduce LD in hPSCs. Interestingly, a study showed that NVB treatment did not improve HDR in human primary T cells ^35^. We note that the treatment regime and cell type used therein are different from this study, suggesting the choice of DNA repair pathway may be complex and context dependent. Previous studies showed that POLQ deficiency does not affect the genetic stability and development of *P*.*patens* ^36^. Thus, small-molecule inhibition of POLQ and/or delivery of recombinant RPA offers a simple, convenient, and potentially safe way to reduce the risk of the unwanted LD and improve the HDR efficiency in hPSCs.

## METHODS

### Cell culture

H1 hESC line was purchased from WiCell Institute. H1-iCas9 ESC line is a gift from Danwei Huangfu’s lab. The wild-type iPSC line was reprogramed and well characterized in previous studies ^37,38^. The study was approved by the KAUST Institutional Biosafety and Bioethics Committee (IBEC). All cell lines were cultured in Essential 8 medium (ThermoFisher, Cat# A1517001) in rhLaminin-521 (ThermoFisher, Cat# A29249) coated wells with medium change daily.

### Plasmids and lentiviral packaging

Oligonucleotides containing gRNA sequence were Annealed and later cloned into lentiGuide-puro plasmid (Addgene Cat # 52963) followed by the published protocol ^39^. The full-length RPA including RPA1, RPA2, RPA3 open reading frames (ORF) were cloned from cDNA of H1 ESCs and GFP ORF was cloned from pInducer21 (Addgene, Cat # #46948). Subsequently, the ORFs of RPA and GFP were inserted into pInducer21 using Gateway cloning method. The sequences were confirmed by Sanger sequencing. The gRNA lentiGuide-puro, newly constructed vectors and pEGIP*35 (Addgene, Cat #26776) were packaged into lentivirus individually. Briefly, the plasmid was premixed with packaging vectors, then transfected into HEK293T using lipofectamine 3000. The lentivirus was harvested two times after 48 hours and 72 hours. The lentivirus was concentrated by PEG-it Virus Precipitation Solution (System Biosciences) and stored in –80°C freezer.

### siRNA transfection

The protocol of esiRNA transfection was adapted to the instruction of lipofectamine RNAiMAX reagent (ThermoFisher, Cat# 13778150). H1-iCas9 cells were harvested after 1 hour 10 µM Rocki treatment. The esiRNA/RNAiMAX solution was prepared for 3 wells per siRNA of 12-well format plate as the recipe: Mix 1 was prepared by adding 13.5 µl RNAiMAX reagent into 225 µl opti-MEM and vertexing for few seconds. Mix 2 was prepared by adding 90 pmol esiRNA into 225 µl opti-MEM + 90 pmol and pipetting few times. The esiRNA/RNAiMAX solution was done by adding Mix2 into Mix1 and incubating for 5 min, which was used for resuspending 1.5 million cell pellet. After 30 min incubation with esiRNA/RNAiMAX solution, the cells were aliquoted equally into 3 rhLaminin-521 coated wells and cultured in 37°C, 5% CO2 incubator. The cell samples were collected after 24 for knockdown efficiency analysis.

### Quantification PCR (qPCR)

The RNA was extracted by RNeasy Mini kit (Qiagen, Cat #74106) and reversed transcribed to cDNA using iScript Reverse Transcription Supermix (BioRad, Cat# 1708840). The qPCR was performed on a CFX384 real-time PCR detection system (BioRad) using SsoAdvanced Universal SYBR Green Supermix (BioRad, Cat# 725270).

### Flow Cytometry

For PIGA gene edited samples, the gRNA lentivirus infected H1-iCas9 cells were treated with 2 µg/ml doxycycline for 2 days to induce Cas9 expression for gene editing. After the doxycycline treatment for 10 days, the cells were harvested and washed twice by PBS buffer containing 3% BSA and filtered through 70 µm strainer. For each sample, 100, 000 cells were stained with 2 µl FLAER Alexa488 (Cederlane, Cat# NC9870611) in 100 µl PBS buffer containing 3% BSA for 15 min at room temperature. The stained cells were washed once PBS buffer containing 3% BSA. For GFPmut correction samples, the cells were harvested after 3 days post-electroporation and pass through 70 µm strainer. The cells were resuspended in 200 µl FACS buffer containing 1 µg/ml DAPI and loaded on a FACS Aria II cytometer for analysis.

### CRISPR-Cas9 genome editing

WT and Hifi-Cas9 were purchased from IDT. The gRNAs used in this study were designed using Benchling (https://www.benchling.com/crispr) www.benchling.com/crispr) and their sequences were shown in Table 1. The gRNAs were obtained either through in invtro transcribed by MEGAshortscript™ T7 Transcription kit (ThermoFisher, Cat#AM1354) or ordered through IDT as Alt-R crRNAs or sgRNAs. For each electroporation, 50 pmol of Alt-R gRNA and 50 pmol of Cas9 were mixed and incubated at room temperature for 10 mins to form ribonucleoprotein (RNP). Buffer R (from the Neon system kit) was added into RNP to make 10 µl final volume. 200,000 single cells were electroporated using Neon system (ThermoFisher) with the setting of 1600 V, 10 ms width and 3 pulses. For HDR study, 30 pmol ssODN and different amount RPA were added into 50 pmol RNP and electroporated into pEGIP*35 HPSCs. The cells were seeded in one well of 24-well plates immediately after electroporation.

### PacBio and Nanopore sequencing

Genomic DNA of edited cells was extracted using Blood & Tissue Kit (Qiagen, Cat# 69506). The UMI labeling was performed followed the published protocol ^7^. Briefly, the target locus was labeled by one-cycle PCR using UMI primer (table 1) in a 25 µl PCR reaction including 50 ng genomic DNA, 1 µM UMI primer, 12.5 µl 2X Platinum SuperFi PCR Master Mix (ThermoFisher, Cat# 12358010), following the program: initial denaturation at 98 °C for 70s, gradient annealing from 70 °C to 65 °C with 1 °C/5 s ramp rate, extension at 72 °C for 7 min, and hold at 4 °C. The UMI labeled DNA was purified by 0.8x AMPure XP beads, then mixed with universal primer and PrimeSTAR GXL DNA polymerase (Takara, Cat# R050A) and amplified following the program: initial denaturation at 95 °C for 2 min, 98 °C for 10 s, 68 °C for 7 min for 30 cycles, 68 °C for 5min, and hold at 4 °C. The amplicon was used for PacBio or Nanopore library preparation.

For Nanopore sequencing, the library preparation was done using the ligation sequencing kit (Oxford Nanopore Technologies, Cat# SQK-LSK109) followed its standard protocol. The Nanopore sequencing was performed on an Oxford Nanopore MinION sequencer using R9.4.1 flow cells. The reads were base called using Guppy basecaller (v5.0.7). Library preparations of PacBio sequencing were performed with the Sequel Sequencing Kit 3.0, and loaded on the PacBio Sequel instrument with SMRT Cell 1M v3 LR Tray. PacBio official tool termed ccs (v3.4.1) was used to generate HiFi Reads. All procedures were preformed according to manufacturer’s protocols.

Data analysis was performed using VAULT as described previously ^7^. In brief, the UMI primer sequence, fastq file and reference amplicon sequence were provided to the algorithm. VAULT will extract mappable reads followed by extraction of UMI sequences from reads. Reads will then be grouped based on their UMI sequences, and used for parallel analysis of SNVs and SVs. The *vault summarize* command was used to generate the analysis summary.

### Recombinant human RPA protein

Human RPA was expressed and purified as described previously ^40,41^. Briefly, the cloned plasmid was transformed into BL21 (DE3) E. coli. The cells were grown in 2YT media at 37 °C to an OD_600_ of 0.7 and protein expression was induced with 0.5 mM IPTG and further incubated for 4-6 hr at 37 °C. The cells were collected by centrifugation and lysed by lysozyme and sonication. The supernatant was loaded onto HisTrap HP 5 ml column (Cytiva) followed by HiTrap Blue affinity column (Cytiva). RPA fractions containing all subunits were concentrated and loaded onto HiLoad 16/600 Superdex 200 pg column (Cytiva). RPA protein fractions were flash-frozen and stored at-80 °C.

### Statistical Analysis

The data in the figures are shown as mean ± SD unless indicated otherwise. Comparisons were performed with two-sided Student’s t-test unless indicated otherwise.

## ACKNOWLEDGEMENTS

We thank members of Li laboratory, Professor Wolfgang Fischle and Ms. Fernanda V. Romero for valuable discussion; J. Xu for her administrative support. We also thank BIOPIC core facility at Peking university for PacBio sequencing support. This work was supported by KAUST Office of Sponsored Research (OSR) under Award No. BAS/1/1080-01.

## AUTHOR CONTRIBUTIONS

M.L. and B.Y. conceived the project and designed the experiments. B.Y., Y.J., K.A. performed the experiments. C.B. and J.W. carried out the PacBio sequencing and bioinformatic analysis. M.T. and G.Y. contributed to the biochemistry experiments. B.Y., C.B. and M.L. did the data analysis. S.H., Y.H. contributed to the project design. B.Y., C.B., and M.L. wrote the paper.

## SUPPLEMENTARY INFORMATION

**Fig. s1.**
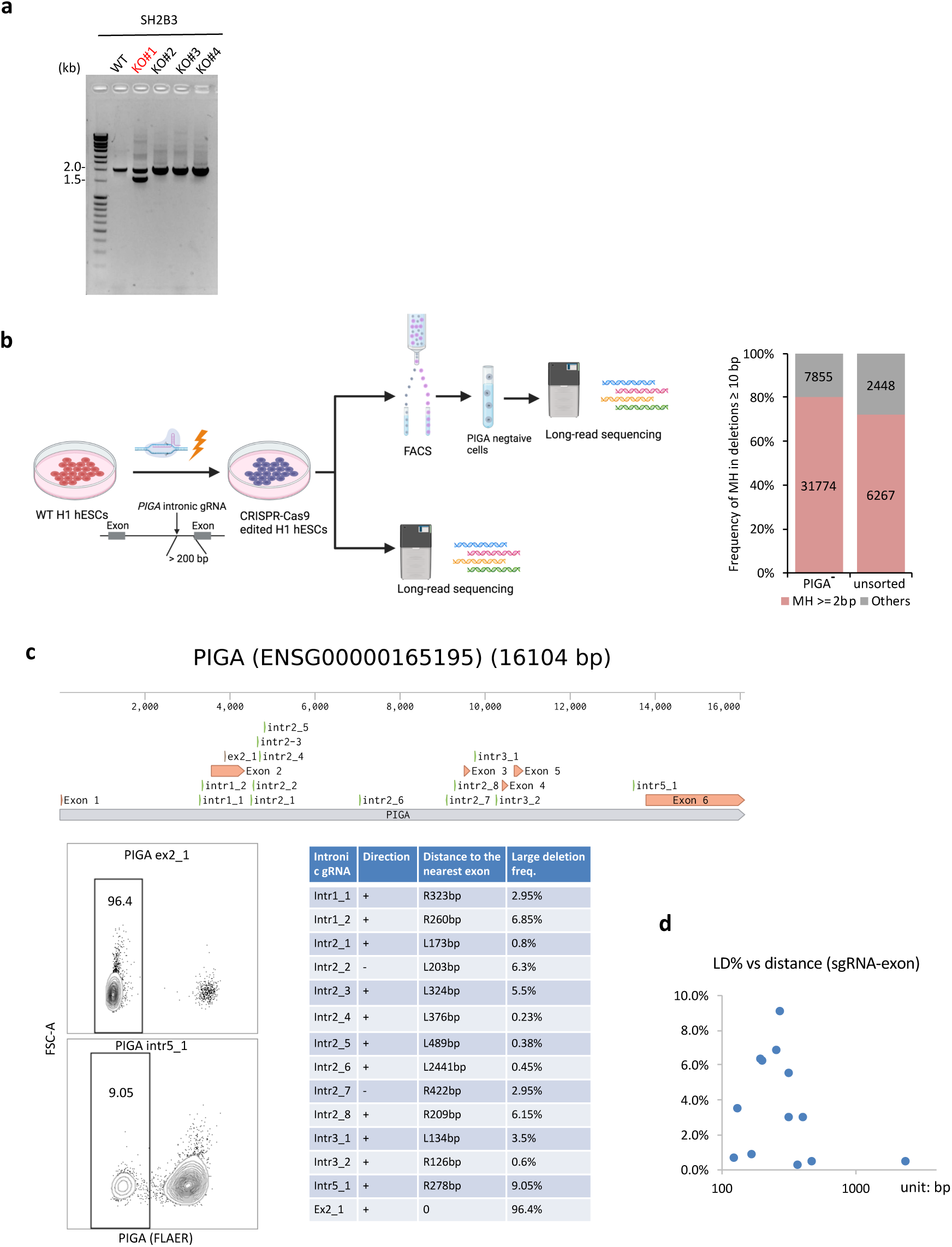
CRISPR-Cas9 genome editing induced LD in human pluripotent stem cells. **a** Agarose gel electrophoresis of long-range PCR products of SH2B3 clones; the red colored sample indicates LD clone. **b** Left: schematic of the strategy to analyze CRISPR-induced LDs in *PIGA* locus; right: MH frequency in deletions ≥ 10 bp quantified from long-read sequencing data. **c** Top: the location of PIGA gRNAs in the PIGA genomic locus; bottom left: example flow cytometry analysis of PIGA expression using the FLAER assay; bottom right: LD frequency of PIGA gRNAs screened, biological replicates *n* =3. **d** Plot of the LD frequency and the distance between the sgRNA and its nearest exon.

**Fig. s2.**
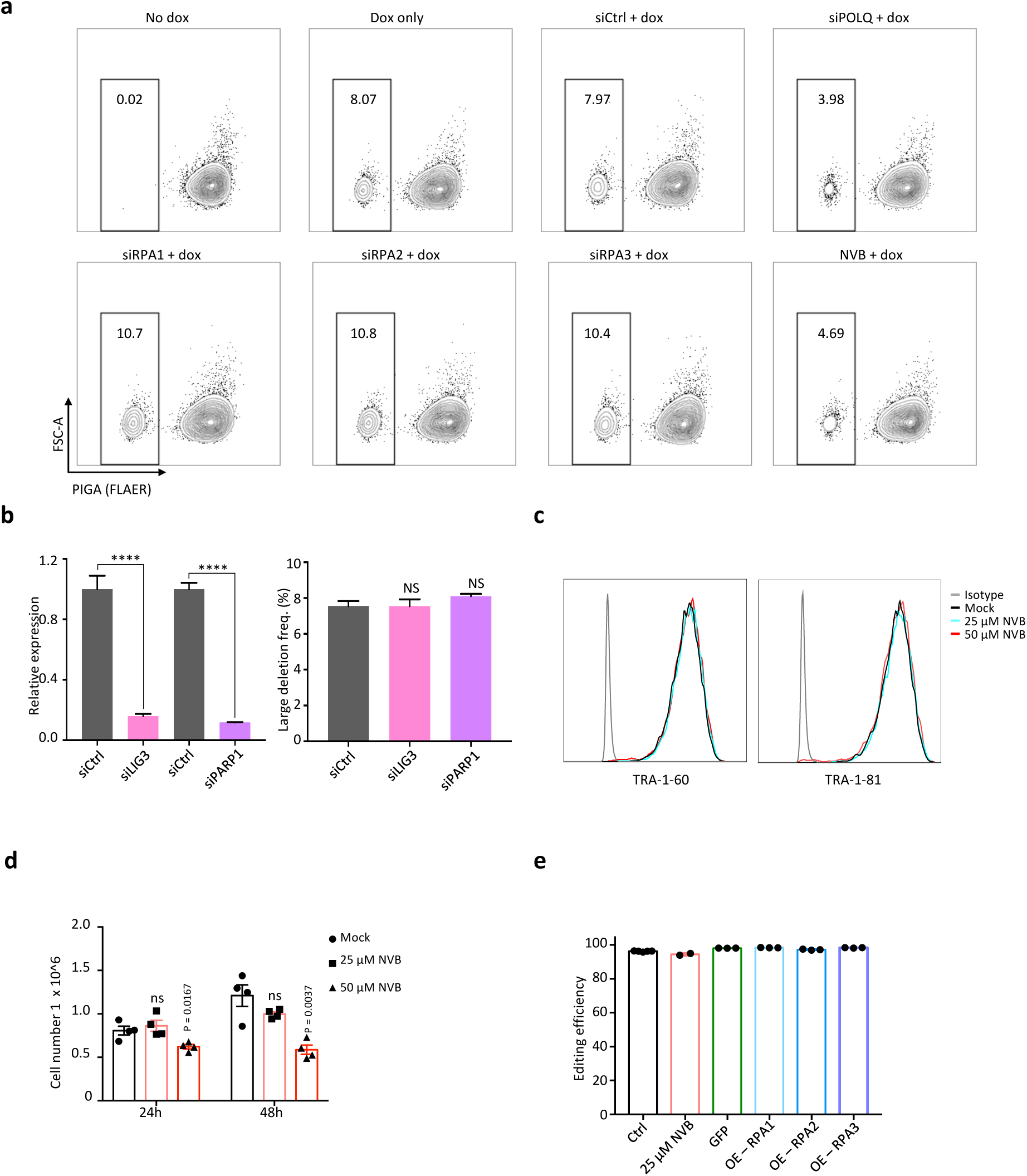
Modulation of MMEJ can regulate CRISPR-induced LD frequency. **a** Flow cytometry analysis of PIGA expression. **b** Left: relative mRNA level of siRNA target genes, LIG3 and PARP1; right LD frequency quantified by FACS, biological replicates *n* =3. **c** FACS analysis for pluripotency markers in NVB treated H1 hESCs. **d** Live cell count of H1 hESCs at 24 hr and 48 hr after NVB treatment, biological replicates *n*=3, ns: not significant. **e** CRISPR-Cas9 editing efficiency using an exonic PIGA guide RNA, ex2_1 sgRNA, quantified by FACS, biological replicates *n*=3.

**Fig. s3.**
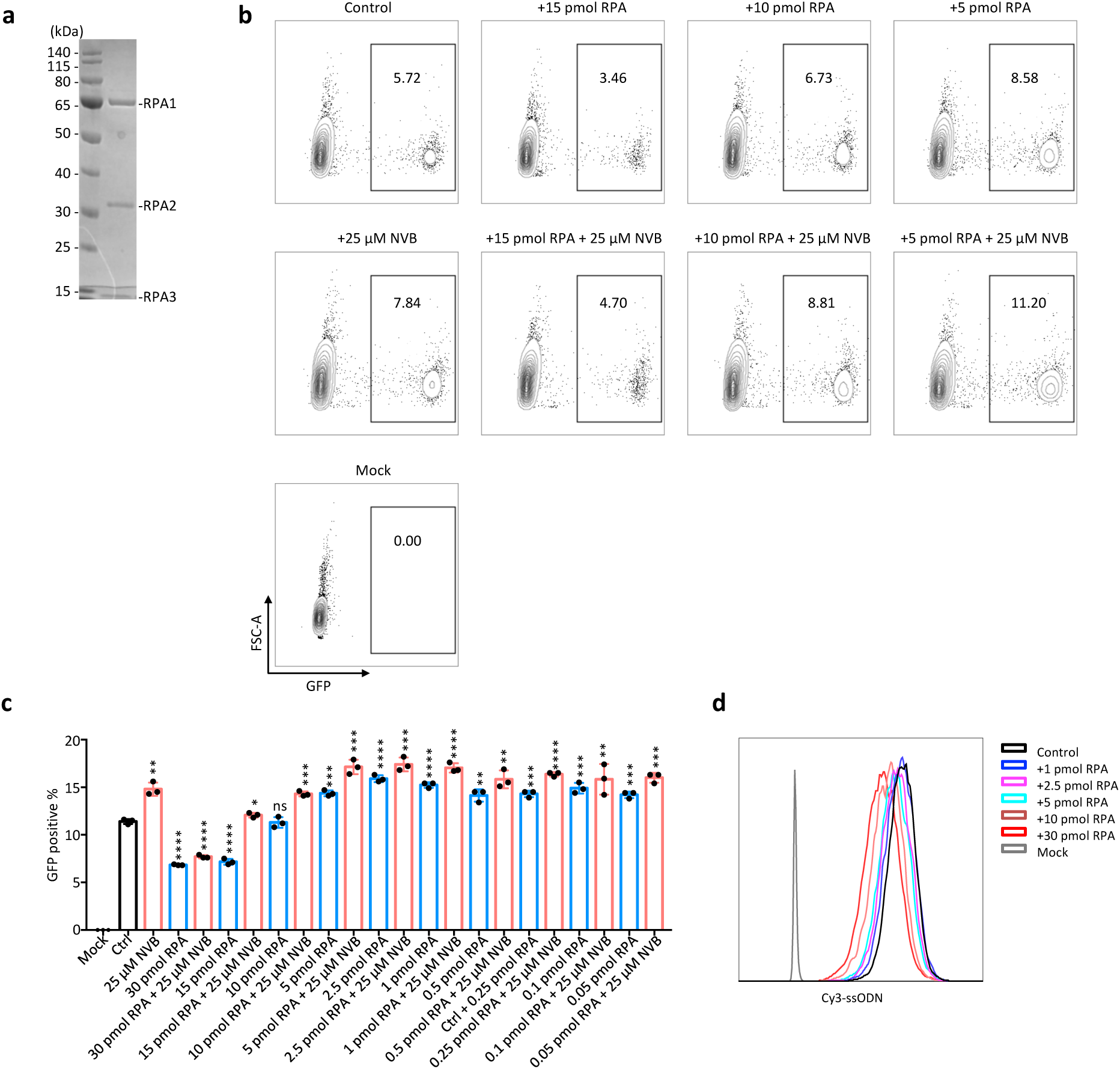
Modulation of MMEJ regulates CRISPR-induced LD frequency. **a** Representative Coomassie-stained SDS-PAGE images of RPA protein complex including the RPA1, RPA2 and RPA3 subunits, kDa: kilodalton **b** Flow cytometry analysis of GFP expression. **c** GFP positive population quantified by FACS, biological replicates *n* =3; * *p* < 0.05, ** *p* < 0.01, *** *p* < 0.001, **** *p* < 0.0001, ns: not significant. **d** Delivery efficiency of Cy3-ssODN mixed with recombinant RPA.

